# Cellular logics bringing the symmetry breaking in spiral nucleation revealed by trans-scale imaging

**DOI:** 10.1101/2020.06.29.176891

**Authors:** Taishi Kakizuka, Yusuke Hara, Yusaku Ohta, Asuka Mukai, Aya Ichiraku, Yoshiyuki Arai, Taro Ichimura, Takeharu Nagai, Kazuki Horikawa

## Abstract

The spiral wave is a commonly observed spatio-temporal order in diverse signal relaying systems. Although properties of generated spirals have been well studied, the mechanisms for their spontaneous generation in living systems remain elusive. By the newly developed imaging system for trans-scale observation of the intercellular communication among ∼130,000 cells of social amoeba, we investigated the onset dynamics of cAMP signaling and identified mechanisms for the self-organization of the spiral wave at three distinct scalings: At the population-level, the structured heterogeneity of excitability fragments traveling waves at its high/low boundary, that becomes the generic source of the spiral wave. At the cell-level, both the pacemaking leaders and pulse-amplifying followers regulate the heterogeneous growth of the excitability. At the intermediate-scale, the essence of the spontaneous wave fragmentation is the asymmetric positioning of the pacemakers in the high-excitability territories, whose critical controls are operated by a small number of cells, pulse counts, and pulse amounts.

## Introduction

The spiral wave is spatiotemporal order commonly observed in the diverse range of excitable media^1^, including the chemical^2^ and biological systems^3,4^. In the biological context, the intercellular relay of communication signaling in the form of a rotating spiral is often associated with pathological cases such as neuronal epilepsy^5^, progressive dermal inflammation^6,7^, and ventricular fibrillation^8,9^. The spiral core, also known as the spatial phase singularity (PS), is a self-sustaining structure that is highly robust to external perturbations; a deeper understanding of the properties and generation of spiral waves is needed to manage and regulate the diseases associated with spiral waves.

While the properties of the mature spirals have been well documented^1^, the onset mechanism, especially in the spontaneous spiral nucleation, is still poorly understood. Conceptually, the spiral wave arises from a pair of open ends of fragmented waves, but the fragmentation of the excitation waves never develops spontaneously in homogeneous systems^1^. Thus, experimental induction of the spirals in the homogeneous system requires designed conditions such as a mechanical break in the waves^2^, geometrical constraints^10^, or additional wave initiation at the vulnerable region in the preceding wave^8,11^ to bring the symmetry breaking, i.e., conversion from closed ring waves (symmetric) to fragmented waves (asymmetric). Alternatively, the systems implemented with heterogeneity in their excitability^12,13^, refractoriness^14,15^, or coupling strength^16^ are capable of generating wave fragments. It is still unclear what types of cellular rationales organize these heterogeneities during the spontaneous spiral nucleation in living systems.

The social amoeba, *Dictyostelium discoideum* (*D. discoideum*), is an ideal model of the self-organized pattern formation. During its development, initiated by the nutrient starvation, 10^3-6^ cells establish spiral-shaped aggregation streams with sub-to few-millimeter wavelengths^17^. The intercellular relay of the chemo-attractant cyclic adenosine monophosphate (cAMP) drives the process^18,19^, whose reaction-diffusion dynamics are explained by a simple combination of the local synthesis, degradation, and diffusion^20-22^. The system is consisted of typical excitable cells having three distinct states including the absolute- or relative-refractory and the excited state, wherein the above-threshold input of extracellular cAMP at the relative refractory state induces the transition to the excited state. The uniqueness of this living excitable system lies in its ability to self-organize the wave patterns even from a quiescent initial state, during which the low-to-high transition of excitability is believed to play a role^22,23^. The excitability is the cellular ability to relay cAMP pulses that are positively regulated by repeatedly relaying cAMP pulses, which is realized by a gradual increase of the gene expression controlling the cAMP synthesis, release, and degradation^18,24,25^. Thus, the question is how the symmetry breaking in the form of the wave initiation and spontaneous fragmentation takes place in the growing excitability. This should be ideally addressed by sensitive and large-scale observations of the cAMP pulse dynamics. The realization of such observations, however, has been challenging due to the technical limitations in fulfilling all of the following conditions: 1) high enough sensitivity to detect faint cAMP pulses even in the low-excitability regime, 2) a large enough observation field to capture the spiral cores at the millimeter-scale, and 3) high enough spatial resolution to identify the pacemaking (PM) cells whose presence is expected to be rare.

In this study, we have developed an improved imaging strategy that allows the bird’s eye view analysis of the cAMP pulse dynamics with a field coverage of > cm^2^ at the single-cell resolution, and have found that the spontaneous wave fragmentation develops in close association with the heterogeneously structured excitability that is highlighted by the fine-mapping of the cumulative pulse count. The analysis at single-cell resolution also identified the critical phenomena in the temporal evolution of local signaling dynamics at the center of the highly pulsing areas, of which a small number of cells controls the collective behavior. Along with the modeled dynamics, we emphasize the functional importance of the microscopic heterogeneity in the initial excitability, which makes the cellular discreteness as the origin of the symmetry breaking.

## Results

### Trans-scale imaging of intercellular communications of 130,000 cells

To analyze the onset dynamics of the spiral nucleation, cellular cAMP dynamics were detected by using a fluorescent reporter Red-FL2, a fusion protein of cAMP indicator Flamindo2 and mRFPmars, whose ratiometric imaging allows quantitative analysis free from motion artifacts (**Fig. 1a**). Instead by using conventional microscopes, exceptionally largescale imaging was achieved by the newly developed imaging system. For short, the system is featured by an objective lens with a large diameter (∼ 50 mm, magnification = 2, N.A. = 0.12), a hundred megapixel CMOS image sensor with a small pixel size (2.2 µm), that eventually allowed the snapshot of 130,000 cell / 14.6 × 10.1 mm^2^ with a spatial resolution of 2.3 µm (**Fig. 1a, 1b**, the detail of the imaging system is described elsewhere). To minimize any damages on our sample associated with live imaging, cAMP dynamics were imaged every 30 sec, being high enough temporal resolution to dissect the timing of cellular cAMP pulse whose FWHM (full width at half maximum) is ∼90 sec at 6-15 min interval (**Fig. 1c**). After 12 hrs of observation started at 4 hrs-post-starvation, we obtained ∼360 G-bites of image sequences that were spatially discretized into 12,236 regions of interest (ROIs, 100 ×100 pxl each containing ∼ 100 cells) and were digitally processed for the analysis of the wave dynamics. The primary focuses of the analysis were the wavefront, oscillation phase, recovery state, and pulse counts (**Fig. 1d, Supplementary Fig. S1**).

**Figure 1.**
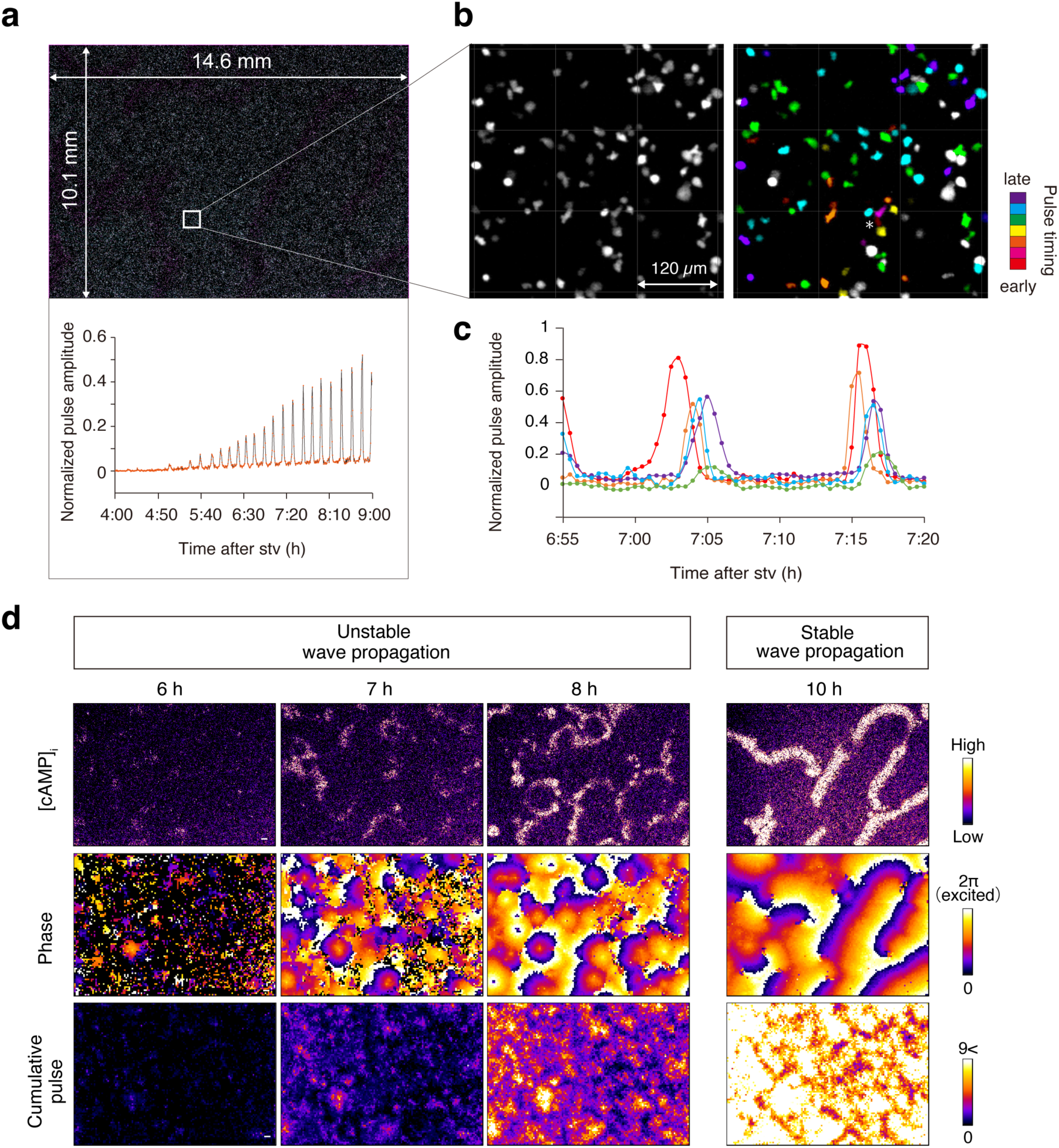
Large-scale Imaging of cAMP Pulse with 1-cell Resolution. (**a**) One-shot-imaging of ∼130,000 cells with 1-cell resolution (top). cAMP pulse detected by the ratiometry of mRFPmars and Flamindo2 for a population (bottom). Average of ∼100 cell data in the white box). (**b**) Close-up view of box in (a), color-coded by the pulse timing. (**c**) Representative cAMP pulses for cells in (b). (**d**) Still images of [cAMP]_i_, oscillation phase, and cumulative pulse counts. Phenotypic features in this culture condition were shown on the top. See also **Supplementary Fig. S1**. Scale bars, 0.5 mm.

### The spontaneous fragmentation of the cAMP waves

To find spatial locations of the wave fragmentation, we reconstructed the phase maps from the temporal dynamics of the cAMP pulses in each locality (**Supplementary Fig. S1**). Because the PS is the singular space in which the entire oscillation phase is encircled, the emergence of the PS in the phase map represents *de novo* wave fragmentation. The localized pulse in two ROIs (asterisk in **Fig. 2a**, 7:32) propagated outward as a symmetric, closed ring, and was broke into the wave fragment whose open ends were marked with a pair of PSs (filled circles in **Fig. 2a**, 7:37). While a part of fragment extending from the counter-clockwise rotating PS (cyan circle in **Fig. 2a**, 7:38) was disappeared through the annihilation with incoming PS (dashed magenta in **Fig. 2a**, 7:38) from right, another fragment from the clockwise rotating PS (magenta) showed a long survival for more than the next 60 min.

**Figure 2.**
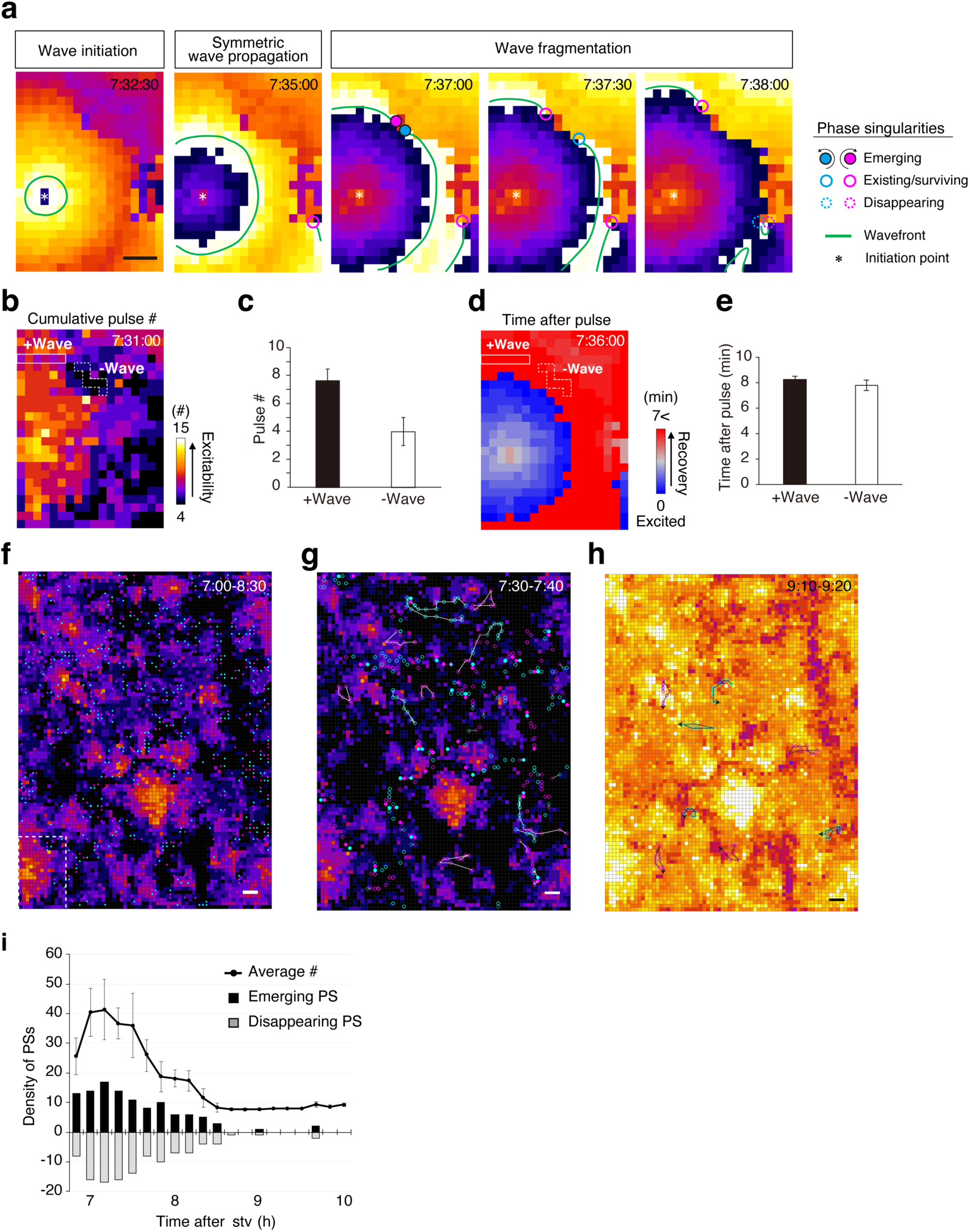
Spontaneous Fragmentation of cAMP Wave. (a) Phase representation of the wave propagation. Green traces represent the wavefront. The asterisk at 7:32 (post starvation) is the point source of the outward propagating wave. A pair of rotating PSs emerges at 7:37. (**b–e**) Comparison of the local property in the wave-permissive (solid) and -blocking regions (dashed border). The cumulative pulse counts (b and c) and the time after pulse (d and e) are measures of the excitability and recovery state, respectively. Error bars represent s.d. (**f**) Spatial distribution of all the emerging PSs during 7:00–8:30 imposed on the map of the cumulative pulse counts at 7:45 of the development. (**g** and **h**) Trajectories of PSs during 7:30–7:40 (g) and 9:10–9:20 (h) imposed on the map of the cumulative pulse counts at 7:30 and 9:10 of the development, respectively. Maximum, minimum, and average of the cumulative pulse counts are 15, 0, and 5.50±2.15 (s.d.), respectively, for (g) and 25, 8, and 14.71±2.12 (s.d.), respectively, for (h). (**i**) Temporal change in the number of PSs. The line graph is the average density (#/whole view) in 10 min data. The histogram presents the numbers of emerging (black) and disappearing (gray) PSs. Error bars represent s.d. Scale bars are 0.5 mm.

What makes propagating waves fragmented in this living excitable system? For the simplest expectation, the low excitability and/or low recovery in the refractoriness would locally block the wave propagation, thus causing the circular wavefront to break open. We examined the properties of the signal-receiving region for the presence or absence of the wave propagation and investigated which parameters were low specifically in the wave-blocking region (**Figs. 2b–2e**). The cumulative number of pulses, *i*.*e*., a measure of the excitability (**Supplementary Fig. S2; Supplementary Note S1**), in the wave-blocking region was found to be approximately one-half of that in the wave-permissive region (**Figs. 2b, 2c**). The elapsed time from the previous excitation in the wave-blocking region, *i*.*e*., an indication of the recovery state, was almost identical to that in the wave-permissive region (**Figs. 2d, 2e**), suggesting that the local difference in the excitability but not recovery state causes the fragmentation of the traveling waves. The importance of the spatially different excitability was further supported by a quantitative analysis of the spatiotemporal dynamics of the PSs in the development (7 to 10 hours of development). The spatial distributions of all the emerging PSs (n = 738) were localized at the boundary between the high- and low-pulsing regions in the heterogeneously structured excitability that was highlighted by the high-resolution map of the cumulative pulse counts (**Fig. 2f**). To further understand the relationship between the PSs and structured excitability, trajectories of the PSs were investigated. As a result, we found that the spatially dynamic trajectories of the PSs in the early development (**Fig. 2g**, 7:30) were localized at the edges of the highly pulsing islands, while those in the regions with the homogeneously high excitability at the later development (**Fig. 2h**, 9:10) were less mobile. The presence of a critical period for the spiral nucleation can explain the temporally different trajectories of the PSs. The developmental changes in the PS density presented a bell-shaped curve with the peak at 7:10 of the development, and both the appearing and disappearing PSs were limited to the early development (until 8.5 hours of the development, **Fig. 2i**). These observations suggested that the transiently structured excitability in the critical period played an important role in the spontaneous wave fragmentation.

The functional importance of the macroscopically structured excitability in the spiral nucleation was tested by the perturbation experiments focusing on the oscillation phases and the landscape of the excitability (**Fig. 3**). The phase resetting was done by a bath application of cAMP and the spatial resetting was newly introduced as follows; the *D. discoideum* cells can be easily detached from a culture dish by intensive pipetting, then the cells restart the cAMP pulse dynamics after they settle on the dish with the cellular positions being randomized (**Supplementary Note S2, Supplementary Fig. S3**). The phase resetting at the post-critical period diminished the existing spirals allowing only the reappearance of the concentric waves (**Figs. 3a, 3c**) as had been demonstrated previously^28, 31^. The phase resetting at the critical period, however, is not effective as we observed >50 PSs to have appeared, indicating that cues for spiral cores are temporally specific to the early stage of development and are not affected by the phase resetting. We then tested the spatial resetting at the critical period and observed that the reappearance of PS was reduced to 1/5 of that for the phase resetting (**Figs 3b, 3c**). These results demonstrate that the heterogeneously structured excitability with an mm-size of coherency is the cause of the spontaneous wave fragmentation during the critical period, but not in the post-critical period.

**Figure 3.**
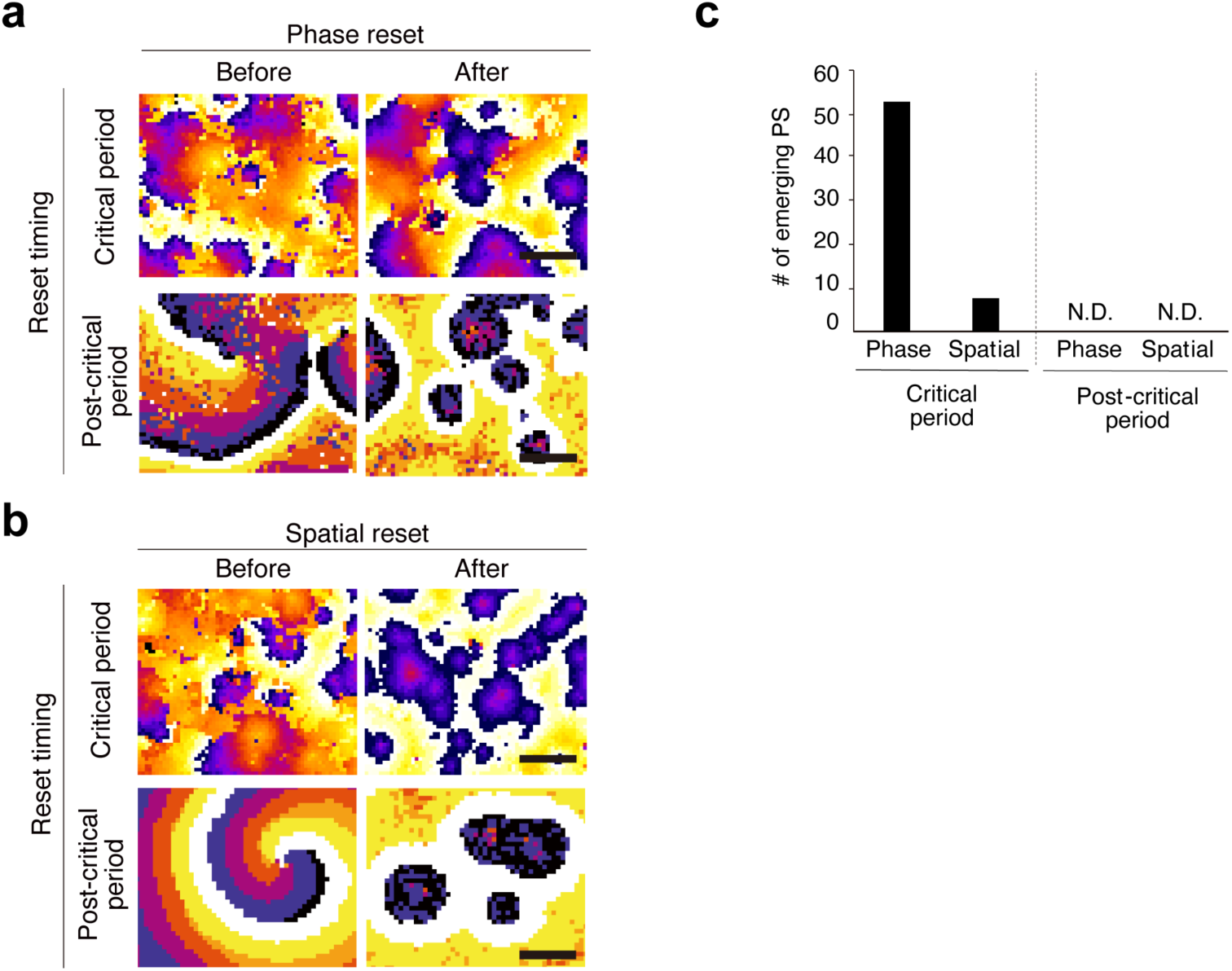
Effects of the Spatial and Phase Resetting on the Spiral Nucleation. (**a** and **b**) Phase images of the wave dynamics before and after the resetting. Phase (a) and spatial resetting performed at the critical period (top) or post-critical period (bottom), respectively. The average number of emerging PSs after the resetting obtained from 2 independent experiments. Scale bars, 2 mm.

In a short summary, to answer the question as to how the propagating waves are fragmented, we have demonstrated that the presence of the heterogeneous excitability and its functionality in the critical period are both essential and that the high/low boundary of the excitability is the cause of the fragmentation in the isotropically traveling wavefront, which subsequently becomes the generic source of the spiral cores.

### The development of the high-excitability territories

How does such spatially heterogeneous excitability develop from the quiescent initial state? The self-organization of the heterogeneity could be explained either by some distinct local rules controlling the pulse-dependent increase in the excitability, or more simply, by different initial conditions such as the spatially biased presence of the PM cells. To examine how the heterogeneous excitability develops, we focused on localities showing the most active or inactive development and analyzed the growth dynamics of the local excitability including the PM activities at cellular resolution. The images of the cumulative pulse counts highlighted the growth pattern of the highly pulsing area (9 neighboring ROIs, magenta box in **Fig. 4a**). The first pulse emerged as early as 4:15 after the starvation, and then the wave-permissive area expanded outward every pulse, each of which originated from inside the highly pulsing territories. In the following development, this area developed to few millimeters in diameter with a pulse count of 9 at its peak (6:30), while the surrounding areas had fewer than 2 pulses (**Fig. 4a**).

**Figure 4.**
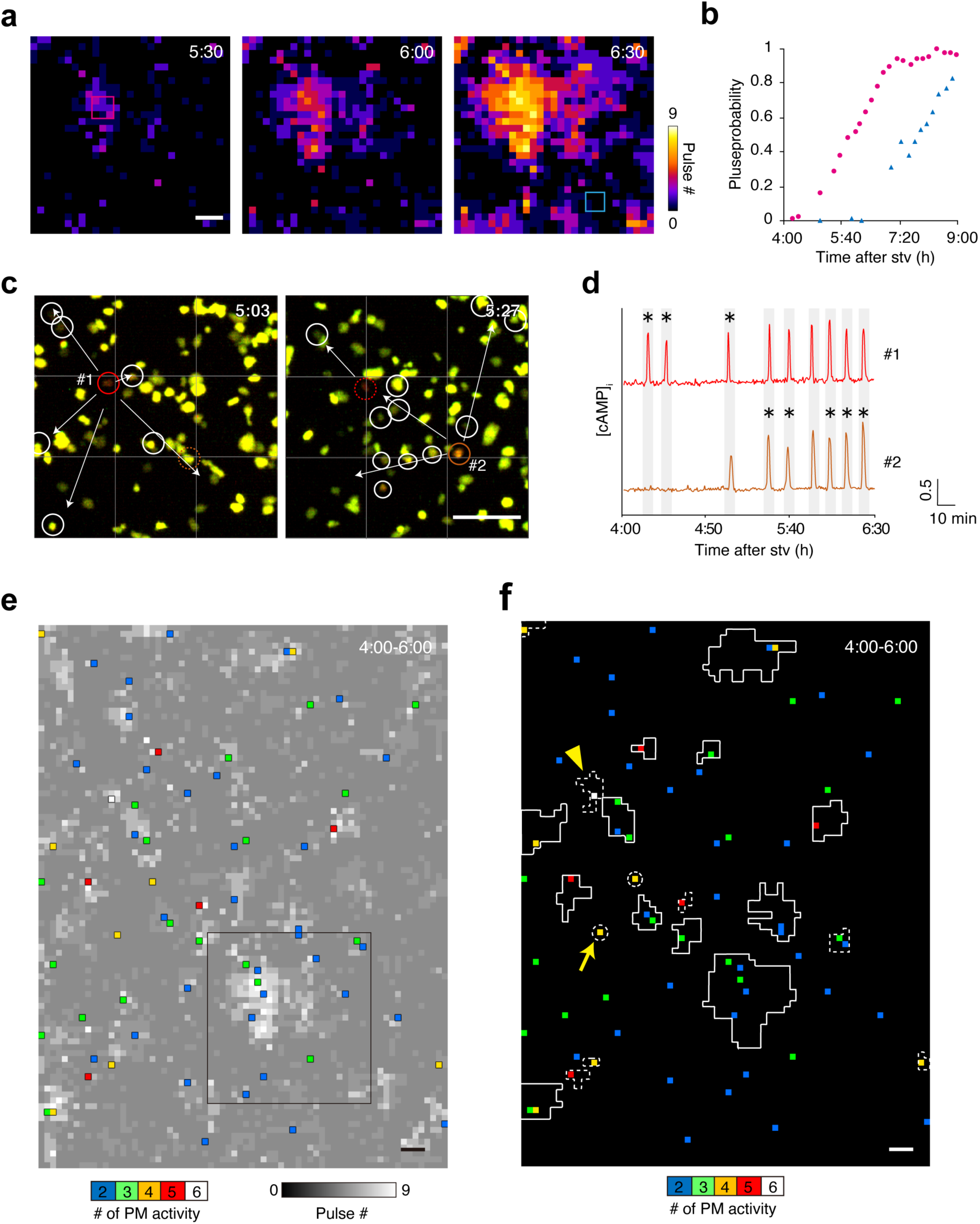
Growth Dynamics of the Highly Pulsing Territory. (**a**) Spatiotemporal growth of the pulse dynamics. (**b**) Pulse probability of the hot spot (9 neighboring ROIs, magenta box in a) and cold spot (cyan box in a) plotted against time. (**c** and **d**) Non-random PM activities of fixed cells. Position (c) and pulse pattern (d) of two PM cells (orange and magenta) and 7 and 12 subsidiary cells for 5:03 and 5:27, respectively (white circles in c) in the hot spot. Temporal flow of the pulse timing, starting from PM cells followed by cells with arrows in this sequence. Grid size is 120 × 120 μm^2^. Asterisks in (d) are the PM pulses. (**e**) Location and activity of PM cells in the analyzed field (∼ 65,000 cells/7.3 × 10.1 mm^2^), imposed on the map of the cumulative pulse counts at 6:00 of the development (gray). Box represents the area analyzed in (a). (**f**) Wave propagated area at 6:00 for representatives of well-(solid) and poor-(dashed) developing territories, imposed on the map of location and activity of PM cells. Scale bars are 0.5 mm.

To examine whether or not the local rules control the pulse-dependent development of the excitability, we quantified the probability of the pulsing cells (number of pulsing cells/total cells in the neighboring 9 ROIs) in two representative localities with high and low pulsing activities (**Figs. 4a, 4b**), hereafter called hot and cold spots, respectively. In the hot spot (magenta in **Fig. 4a**), a cAMP pulse started when 2 out of 107 cells were pulsing (pulse probability of 0.02). As the pulse count increased every 10-20 min, so did the local pulse probability until reaching the plateau (0.95) at 7 hours of the development. In the cold spot (cyan box in **Fig. 4a**, containing 109 cells), the timing of the pulse initiation for cell- and population-level were delayed 1 and 2 hours, respectively, as compared to that of the hot spot. Re-plotting the pulse probabilities for the growing population (> 0.15) against the pulse counts revealed similar growth rates (the half-maximal pulse probabilities yielded the pulse count of ∼5, **Supplementary Fig. S2**). It indicated that the rule controlling the increase in the excitability as a function of accumulating pulses was conserved among different localities. The major difference leading to the distinct timings of the pulse initiation would be the initial condition. Indeed, we found two PM cells showing 8 rounds of leading pulses at the center of the hot spot (inside 3×3 ROIs) together with two supportive PM cells (twice and once activity) around the center for 4.0-6.5 hrs of development (**Figs. 4c, 4d**): No PM cell was present in the cold spot where the population pulse was always triggered by propagating waves from the surrounding regions throughout the development.

### The logics controlling the critical transition

To further investigate the cellular rationales controlling the local development, we asked whether the pacemakers were sufficient to account for the hot spot development, and presumed that the search for the PM activity was important because the PM activities at the single-cell or population level are essential to trigger the wave propagation in the excitable system. Although it has been long believed that the presence of the PM cells is sparse, their actual density and pulse patterns (random or deterministic) have not been analyzed. Our trans-scale observation successfully identified 84 cells (corresponding to 0.13% of the total population distributed in 67 of color-coded ROIs, **Fig. 4e**) which showed the spontaneous and repeated PM activities during the early development (4:00–6:00). While PM activities of these ROIs were evenly distributed along time (data not shown), their position was found to be biased to the center of developing territories as the single or cluster of ROIs (**Fig. 4e**). To more correctly understand the relationship between the pacemaking activity and the hot spot development, we detected the wave permissive area for the latest activity before 6:00. As in the case for the above analyzed hot spot, these highlighted areas for the representative of well-developed territories (solid line in **Fig. 4f**) were found to be associated with ROIs with high PM activities (pulse counts >3). Inversely, when we focused on ROIs with high PM activities (pulse counts >3), several ROIs were found to show poor development (dashed lines in **Fig. 4f**). In the most extreme case, no wave propagation was observed for ROI with 4 rounds of PM activities by the single cell (arrow in **Fig. 4f**). Collectively, these results suggest that PM cells are essential but insufficient for the development of the hot spot.

To address what mechanisms are needed to drive the local development, in addition to PM activities, we focused on a locality showing moderate development (arrowhead in **Fig. 4f**) and investigated the pulse dynamics at the single-cell resolution (**Fig. 5**). The pulse probability remained low during the first five population pulses, then an abrupt leap broke out at the 6^th^ pulse (19/∼90 cells at 5:25, **Fig. 5a**). Other regions including the hot and cold spots also displayed similar leaps, suggesting that this is a general phenomenon. We hypothesized that such leaps, or the critical transitions (CTs), may play an important role in the evolution of the excitability. To gain more insights into the cellular mechanisms of the CT, we functionally categorized the pulsing cells based on the timing of the coupling to the population pulse (before or after the CT) and the transition pattern of the pulse amplitude. It was possible to group the activities into three distinct classes, which would correspond to the different excitability (**Fig. 5b**): The first group was “leader” characterized by PM activities of the cells (**Fig. 5c**). The large amplitude (>0.4 of the normalized amplitude) throughout the development was a notable feature reflecting the highest excitability. The opposite extreme was the group named “citizens” containing the majority of the cells (>90%) that were characterized by a delayed start of the cAMP pulses after the CT. The amplitude of the pulses in this group increased from a small value, suggesting the low excitability in the beginning. The remaining groups were “follower” (6 cells), showed a gradual increase in the pulse amplitude like the citizens, but the onset was advanced to the CT.

**Figure 5.**
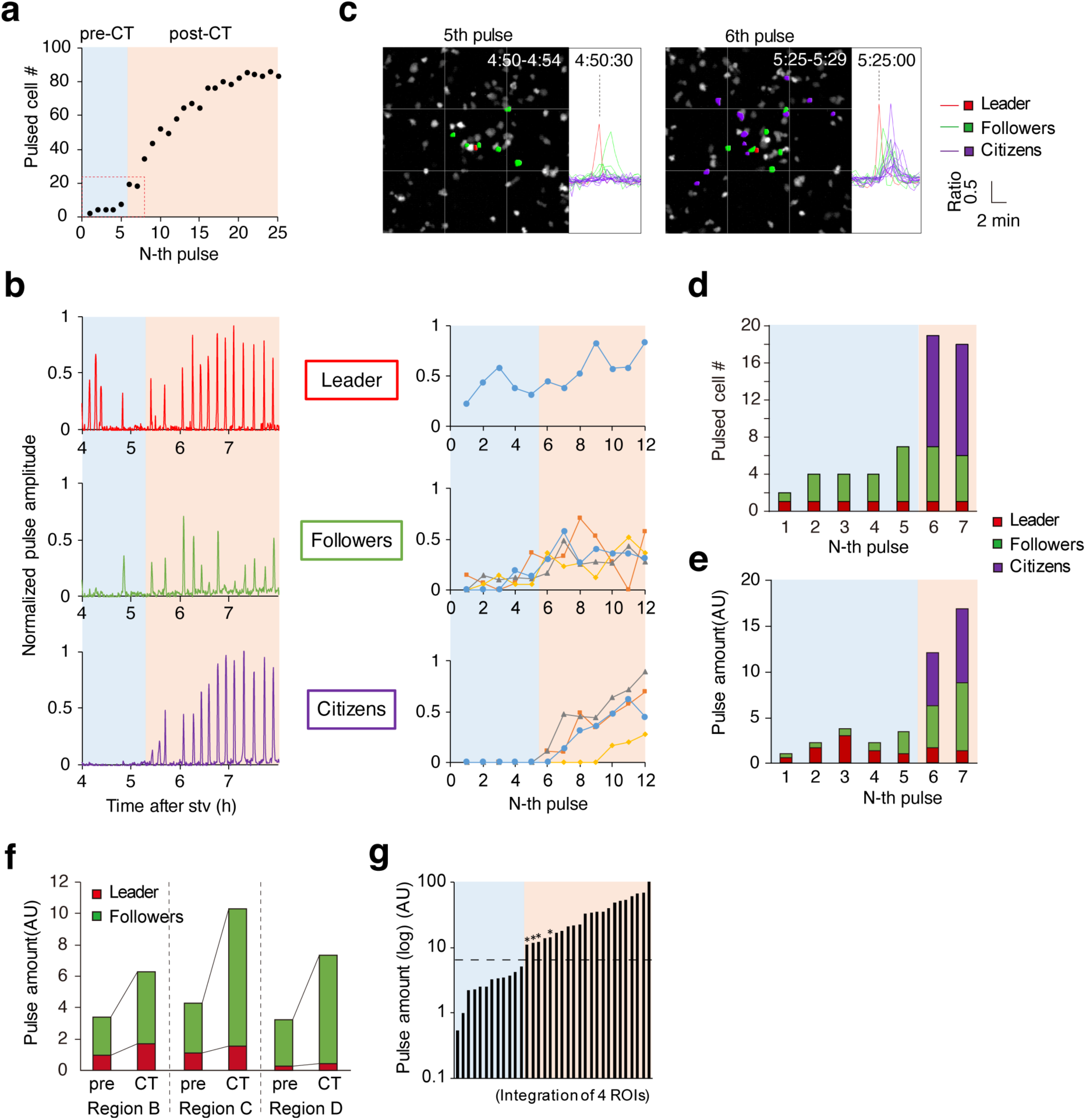
Critical Transition in the Local Pulse Dynamics. (**a**) Burst in the number of pulsing cells for 9 ROIs containing arrowheded PM in Fig. 4f. Pulsed cell number plotted against the number of population pulse. Gray and orange masks discriminate before and after the critical transition. (**b**) Pulse dynamics of leader, follower, and citizen cells. Representative 1-cell pulse and the temporal change of the peak amplitude along with the N-th population pulse. (**c**) Location and the pulse timing by the cell group at pre-CT and CT. Grid is 120 x 120 μm. (**d, e**) The number(d) and quantified cAMP pulse(e) summed by the cell type. (**f**) Changes in the pulse amount of leader and followers in 3 hot spots at pre-CT and CT. (**g**) The ensemble of pulse amount of 4 hot spots before and after the CT, sorted by the magnitude. Asterisk is the pulse amount at CT. The dashed line is the estimated threshold for the CT. See **Supplementary Fig. S1** for the location of the analyzed area.

To unveil the mechanisms leading to the CT, we asked the difference in the number of pulsed cells and signal strengths among three groups between pre-CT and CT. In definition, it is natural that both the number and amount of pulsed citizens were increased at CT (**Figs. 5d, 5e**), but it is not plausible to consider citizens bringing the CT. Since the position and pulse timing of citizens at CT is distal and behind to that of the leader and followers, respectively, indicating that the pulse of citizens was the result of CT rather than the cause. Instead, we observed the increase of pulse amount of followers, while its number was kept constant before and after the CT (**Fig. 5d**). This suggested that the driving force of the CT would be the followers. Supporting this idea, we observed similar increases in the pulse amount of followers in the other two hot spots (**Fig. 5f**). Furthermore, the ensemble of the pulse amount from 4 different hot spots for pre- and post-CT suggested that conserved mechanism for CT, whose threshold-like behavior was controlled by estimated pulse amount, being 4.5 μM(AU) of summed pulse/9ROIs (**Figs. 5f, 5g**).

Taken together, these findings have demonstrated that the development of hot spots requires both PM activities of the leaders and effective amplification by the followers, the latter controls timings of the CT leading to the locally distinct excitability.

### The origin of the symmetry breaking

Although the above results have demonstrated the development of the macro-scale heterogeneity of the excitability, this did not explain the mechanisms of the spontaneous wave fragmentation. Mesoscopically, one of the key factors leading to the wave fragmentation is the off-center positioning of the wave initiation loci in the high-excitability territory, which should originate from microscopic asymmetry. To examine the origin of the symmetry breaking, we performed the numerical simulations based on cellular automata (CA). The genetic feedback model was originally introduced by Levine^23^. The schemes with slightly different assumptions from the first model were later developed by several groups^28,32^. The model with these schemes describes the two-dimensional reaction-diffusion dynamics of the extracellular cAMP, where the spatially heterogeneous excitability evolves through the pulse-dependent positive feedback on the pulsing potential. Although the model of these schemes well explains the spontaneous wave fragmentation in the presence of densely distributed PM cells, we found that this was not the case when the density of the pacemakers was altered so that it was lower than the level observed in real systems (**Figs. 4e, 4f**). The schemes assume the homogeneous initial condition of the excitability (*E*_ini_), which allows the spatially isotropic development of the local excitability. The asymmetric positioning of PM activities in the wave-permissive territory can be realized only when a few of these territories are closely packed to each other. We overcame the inability of the schemes to produce spiral waves at low pacemaker density by introducing the distributed *E*_ini_ instead of the homogeneous one. As shown in **Fig. 6**, the fixed ROI with PM activities yielded the anisotropic growth of the high-excitability territory. Also, the simulation was able to reproduce the spontaneous wave fragmentation in the high-excitability territory (**Figs. 6a, 6b**) as observed in the real system (**Figs. 6c, 6d**). The variations of the size of the high-excitability territories and spiral wave formation for the distributed *E*_ini_ were reproducible (**Figs. 6e, 6f**), whereas the homogeneous excitability was not (**Supplementary Fig. S4**). The reproducibility further confirmed the validity of our assumptions. The pattern of the distribution was not critical since the wave fragmentation itself broke out with a variety of non-homogeneous *E*_ini_, including the long-tailed, random, and uniform distributions (**Supplementary Fig. S4**). Although the *E*_ini_ is not directly measurable in real systems, the good agreement with the observed and simulated wave dynamics in both the unperturbed and perturbed conditions (**Supplementary Fig. S4**) has demonstrated that the distributed *E*_ini_ is the origin of the symmetry breaking in the self-organized spiral nucleation of the excitable system in our settings.

**Figure 6.**
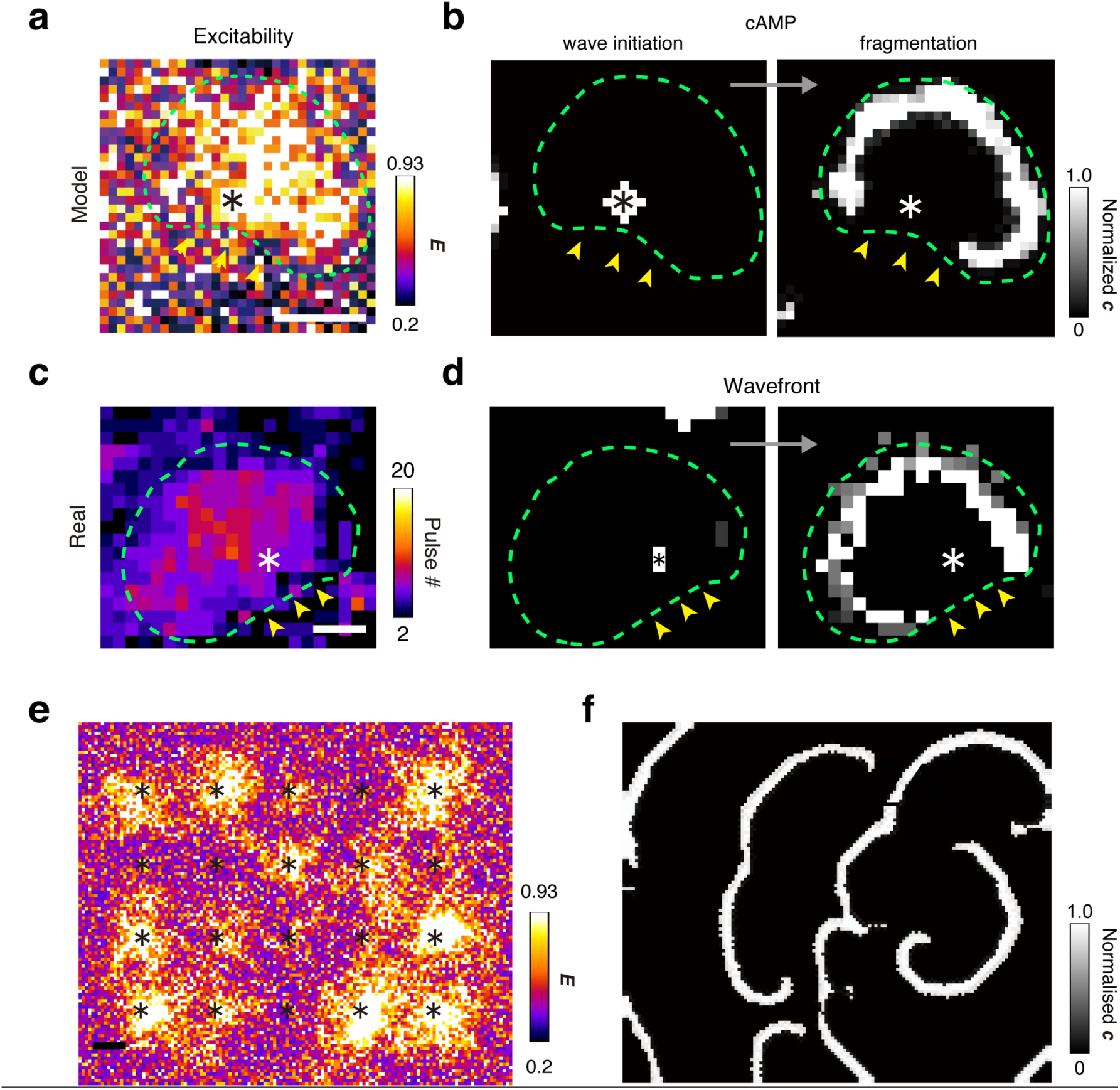
CA Modelling with Few Pacemaker Activities. (**a–d**) Comparison of simulated (a and b) and observed (c and d) development of the single high-excitability territory (a and c) and wave patterns (b and d). Off-center positioning of the fixed pacemaker (asterisk) and high-excitability territory (dashed green line). Arrowheads show the spontaneous wave fragmentation in the single territory. (**e**) High-excitability territories with variable shapes and sizes developed by 20 symmetrically positioned pacemakers in a larger-scale simulation. (**f**) Spontaneous generation of the spiral wave from (e). An exponentially distributed *E*_ini_ (mean value, 0.2) is considered. Asterisks denote the positions of pacemakers. These results were replicated on three different data sets of *E*_ini_. Scale bars, 0.5 mm. See also **Supplementary Fig. S4**.

## Discussion

Our trans-scale analysis leads the three major insights at each scale into the mechanisms of the self-organized spiral nucleation. At the macro-scale, the traveling front of the excitation wave is fragmented at the edge of the heterogeneously structured excitability. The micro-scale analysis has revealed that, for its growth, the involvement of both the PM activities of the spatiotemporally non-random, rare leader cells and effective amplification by the follower cells is crucial. Importantly, these findings themselves do not explain the central mechanism of the symmetry breaking. That is because the key concept of the spontaneous wave fragmentation lies in the off-center positioning of the pacemakers in the high-excitability territory. The meso-scale analysis together with the mathematical modeling has demonstrated that such symmetry breaking is seeded by the non-random activities of the fixed pacemakers. They anisotropically amplify subtle differences in the cellular excitability that eventually cause the spontaneous wave fragmentation autonomous to the high-excitability territory.

We emphasize that, through the use of the trans-scaling imaging techniques for cAMP dynamics, we have unified the mechanisms of the spiral nucleation at distinct scalings (*i*.*e*., micro-meso-macro). Specifically, the sensitive detection of the pulse dynamics even at the low-excitability regime has allowed us to utilize the cumulative number of pulses as an indirect but valuable measure of the local excitability, for which a suitable molecular marker has not been found. The biochemical and genetic studies showed the pulse-dependent increase of the excitability in *D. discoideum* cells^24,25,33,34^. We have also observed similar dependency as summarised in **Supplementary Notes S1**. The increase in both the pulsing ability at the single-cell level and the number of pulsing cells at the population level explains the gradual increase in the amplitude of the population-ensemble cAMP pulses (**Supplementary Fig. S2**). A clear correlation between the oscillation period and the pulse count also supports our hypothesis that the cumulative number of pulses denotes the local excitability (**Supplementary Fig. S2**). The identified geometry of the excitability with local coherency (sub-millimeter-scale) and global heterogeneity (millimeter-scale) is of special significance. This is the first visualization of the spatiotemporal evolution in the growth dynamics of the cellular excitability whose importance is uniquely attributed to the biological self-organization, but not to the chemical models (*i*.*e*., BZ reaction) with non-growing excitability. The anisotropic growth of the high-excitability territory and functional classification of the cells by their distinct excitability have collectively demonstrated the effectiveness of our trans-scale analysis. It should also be effective in the studies of other growing excitable media such as the segmentation clock^35^ and neuronal circuits.

Notably, the so-called “the law of the few” drives the self-organized pattern formation in the discrete system^41^. In each excitable territory, the wave dynamics is initiated by a small number of the PM cells. To our knowledge, this is the first study to estimate the density of such cells (one per 770 cells). The collective behavior of the cell population arises when the critical threshold of ∼4.5 μM of pulses (AU)/∼100 cells/0.13 mm^2^ is reached. For the wave fragmentation, the difference in the effective pulse count between 0 and 10 has functional importance, but not in the count above 10. Such singular properties of the CT governed by the law of the few should apply to the dynamical systems ranging from the cells to the organisms, the eco-systems, and the climates. More efforts in seeing the forest for the trees, or even for the leaves will help to predict and control the system dynamics in the real world^42,43^.

## Methods

### Molecular biology

cDNA encoding mRFPmars-Flamindo2 fusion protein (Red-FL2) whose codon usage was optimized for *D. discoideum* was constructed by using inFusion cloning system (TAKARA), then was cloned into pDM304 and pDM358^44^. Resulting plasmids, pDM304_Red-FL2, and pDM358_Red-FL2, were deposited to the Dicty Stock Center.

### Cell culture

The axenic strain Ax2 was cultured and transformed as described elsewhere^45,46^. For transformation, the cells were washed and suspended with ice-cold EP buffer (6.6 mM KH_2_PO_4_, 2.8 mM Na_2_HPO_4_, 50 mM sucrose, pH 6.4) to 1 × 10^7^ cells/ml. A total of 800 µl of cell suspension mixed with 10 µg of pDM304_Red-FL2 in a 4-mm-width cuvette was subjected to electroporation (two 5 sec separated pulses with 1.0 kV and a 1.0 msec time constant) using a MicroPulser (Bio-Rad). These cells were plated on 4–6 pieces of 90-mm plastic dishes with HL5 medium and incubated at 22°C for 18 hours under the non-selective conditions and then cultured in the presence of 10 µg/ml G418 (Wako). After 4–7 days, colonies with high expression of Red-FL2 were manually picked. Some of these were subsequently transformed with pDM358_Red-FL2, then were cultured in the presence of 35 µg/ml hygromycin (Wako) and 15 µg/ml G418. Clones showing higher expression of Red-FL2 with lower heterogeneity were screened.

### Imaging

The cells expressing Red-FL2 were maintained in HL5 medium on a 90-mm plastic at a density of < 1 × 10^6^ cells/dish. The development was initiated by the 3× washing of the cells with development buffer (5 mM Na_2_HPO_4_, 5 mM KH_2_PO_4_, 1 mM CaCl_2_, 2 mM MgCl_2,_ pH 6.4). These cells were plated on a 35-mm plastic dish at a density of 750 cells/mm^2^. The live-cell imaging was performed by using a custom-built imaging system equipped with the single CMOS image sensor and LEDs illumination (unpublished).

### Numerical simulations

The simulations were performed on a 114 × 136 mesh where *Δ*_x_ = *Δ*_y_ = 0.048 mm (**Figs. 6e, 6f**, and **Supplementary Fig. S4**) using the explicit Euler method at *Δ*_t_ = 0.1 min with synchronous updating in MATLAB. The average of the initial excitability (*E*_ini_) was set to 0.2 for both the homogeneous and distributed cases.

### Equations

To explain the territory-autonomous wave fragmentation, one minor and two major modifications are introduced into the genetic feedback model based on our experimental observations.

Briefly, the genetic feedback model^23,28,32^ is a hybrid cellular automaton (CA) in which the reaction-diffusion dynamics of the extracellular cAMP (*c*) and the pulse-dependent increase in the excitability (*E*) obey the following:

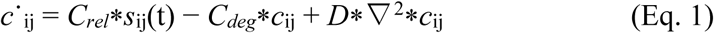

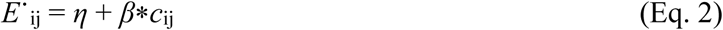

where *C*_*rel*_ is the release rate of cAMP, *C*_*deg*_ is the degradation rate of extracellular cAMP, *D* is the diffusion constant of cAMP, *β* is the feedback strength of the cAMP concentration on the excitability and *η* is the autonomous increase in the excitability over time. Cell_ij_ is treated as a CA whose three discrete states (excited, absolute refractory, and relative refractory) are controlled by the binary operator *s* (*s* = 1 for excited state and *s* = 0 for absolute and relative refractory states). The residence times for the states are *T*_*ex*_ = 1 min, *T*_*abs*_ = 2 min and *T*_*rel*_ = 7 min. Except for the PM activity, a cell receiving an above-threshold cAMP at the relative refractory state is excited where the decreasing threshold *C*_T_ (from *C*_max_ to *C*_min_) obeys the following:

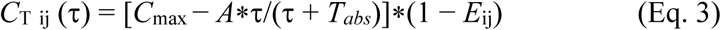

where *A* = (*T*_*rel*_ + *T*_*abs*_) *(*C*_*max*_ − *C*_*min*_)/*T*_*rel*_ and τ measures the elapsed time in the relative refractory state (0 < τ < *T*_*rel*_). After *T*_*rel*_, *C*_T_ is kept to *C*_min_. For the PM activity, a spatially fixed cell is autonomously pulsed depending on the firing probability *p*_fire_, as assumed in ref32.

To reproduce the observed wave dynamics, we modified the above model to

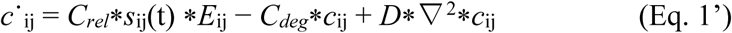

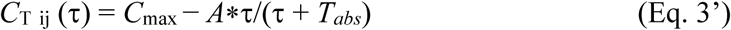

where cAMP is synthesized proportional to *E* (in Eq. 1’). The threshold *C*_T_ is freed from *E* (in Eq. 3’), and *C*_max_ and *C*_min_ are tuned to small enough values to allow the pulse dynamics even at the low-excitability regime. Essentially, these changes do not affect the central dynamics of the model and only increase the dynamic range of *c* and *E*. The major two modifications were introduced to the PM activity and *E*_ini_. The spatial^23, 28^, or temporal^32^ randomness in the PM activity, being the source of the symmetry breaking in the previous schemes, was eliminated by assuming a small number of fixed PM cells (0.6 cells/mm^2^) with almost identical pulse intervals (20 ± 0.45 min). Finally, the distributed *E*_ini_ rather than the homogeneous one was considered, which plays the central role in the symmetry breaking in our modeling scheme. To focus on the functional importance of *E*_ini_, a regular arrangement of the 20 PM cells on the square lattice with a spacing of 1.1 mm was considered.

Collectively, our modifications were minimally considered in (Eq. 1’) and (Eq. 3’). The parameters changed are *C*_max_ = 40 and *C*_min_ = 1, the number of PM cells is 20 (4 × 5 array/5.4 × 6.5 mm^2^) with *p*_fire_ = 0.0005 ± 0.0001, and all the considered *E*_ini_ have a mean value of 0.2, while the other parameters are identical to those of ref. 28. (*E*_max_ = 0.93, *C*_*rel*_ = 300, *C*_*deg*_ = 8 min^-1^, *η* = 0.0005, β = 0.005 and *D* = 2.3 × 10^-7^ cm^2^/sec).

### Image analysis

The ratio images of the background-subtracted and spatially smoothed channels for RFP and Flamindo2 were subjected to the image enhancement of pulsed cells assisted by the supervised machine learning by using AIVIA software (DRVISION tech.). The image field was subdivided into 12,236 ROIs (133 × 92 matrices of 100 × 100 pxl), then the peak detection was performed on time series data for every ROI by using Mathematica (**Supplementary Fig. S1**). The oscillation phase, time after pulse, cumulative pulse counts, and pulse counts were analyzed in each ROI containing ∼10 cells (**Supplementary Fig. S1**). The image reconstruction of these data was performed on the custom-built analysis pipeline using Excel, Mathematica, and Fiji software. For peak detection, the detection sensitivity was tuned to detect > 2 pulsing cell/ROI. After obtaining a peak table for 900 frames of 12,236 ROIs, it was manually corrected to detect 1 pulse cell / ROI with ΔR(Red/FL2) > 7.5 %.

### Quantification of the cAMP pulse at 1-cell

To quantitatively compare the amount of cAMP pulse among cells, we performed a two-step normalization for the emission ratio change of Red/FL2. Considering the identical stoichiometry of red and yellow signals of Red-FL2 among cells, we pre-normalized the baseline ratio values to 1.0. This cancels the different baseline ratio of distantly positioned cells affected by a slight imbalance of illumination strength over the image field. For the pre-normalized ratio data, maximum values at the highest peak were found to reach around 4.1 for >300 cells, that is consistent with the previously reported signal change of Flamindo2 (4-fold). We finally normalized the pulse data of all the examined cells by considering a minimum (1.0) and maximum ratio values (4.1) to 0 and 1, respectively. The sum of the normalized ratios of pulsing cells was utilized to estimate the pulsing activity in a given ROI by using Hill equation: [cAMP]^n^ = *K*_d_*ΔR /(1-ΔR), where Hill coefficient (n) of 1.0 for Flamindo2 (*in vitro*) was employed for a simplicity.

### Statistics

As the distribution of data in this study is not normal, nonparametric statistics were used. All statistical tests were two-tailed. The analysis was performed in the R statistical environment version 3.3.2.

### Data availability

Raw data and codes are available from the corresponding author (K.H.) upon a request.

## Acknowledgments

We thank Y. Yoshihara, J. Kajiwara, K. Morimoto, and other members of the K.H’s laboratory for technical assistance. We also acknowledge the Dicty Stock Center for providing us the pDM304, pDM358, and Ax2 cell lines. This work was supported by a Grant-in-Aid for Scientific Research on Innovative Areas “Singularity Biology (No.8007)” (18H05408 to T.N., 18H05415 to K.H.), a Grant-in-Aid for Scientific Research on Innovative Areas “Spying minority in biological phenomena (No.3306)” (23115003 to T.N., K.H.), the Grant-in-Aid for Young Scientists (A) (18687014 to T.N.), and the Research Program of “Five-star Alliance” in “NJRC Mater. & Dev.” (T.N., K.H).

## Author Contributions

T.K., T.I, and T.N developed the fluorescence imaging apparatus. T.K, A.M., A.I., and K.H. performed experiments. Y.H., T.K., A.M.,Y.A., and K.H. analyzed data. Y.O. and K.H. performed the modeling. Y.H., T.N., T.I. and K.H. designed and conducted the project and wrote the manuscript.

## Supplementary Information

Supplementary Information includes two Supplementary Notes and four Figures.

