## Supplementary Information for "Cellular logics bringing the symmetry breaking in spiral nucleation revealed by trans-scale imaging"

**This PDF file includes:**

Supplementary Notes S1to S2

S1. Cumulative number of pulses as a measure of the excitability

S2. Reset experiments

Supplementary Figures S1 to S4

### Supplementary Notes:

#### S1. Cumulative number of pulses as a measure of the excitability

The population-averaged cAMP pulse shows a gradual increase in its amplitude (**Fig. 1a**), which is explained by the increase in both the pulsing ability at the single-cell level and the number of pulsing cells at the population level. As discussed for **Fig. 5**, the cells are functionally classified into 3 types, and the majority of the total population (>90 %) is the follower and citizen cells, whose pulse amplitudes gradually increase as they experience more numbers of the cAMP pulses (**Fig. 5b**). At the population level, the local pulse probability for the post-critical transition (CT) shows a conserved growth rate among different localities (**Supplementary Fig. S2**), indicating that an increase in the number of pulsing cells also contributes to the gradual increase in the population amplitude (**Fig. 1a**). The oscillation frequency is another point of action for pulse-dependent feedback regulation. Cells that experienced more number of cAMP pulses are expected to show a faster recovery in their refractory processes. Indeed, the spatial pattern of the local excitability and the pulse interval are similar at 8 hours of the development (**Supplementary Fig. S2**), which is quantitatively supported by a clear correlation of these values in 1,672 ROIs (**Supplementary Fig. S2**). Together, these observations validate the working hypothesis that the cumulative number of pulses is a useful measure of the growing excitability.

#### S2. Reset experiments

Perturbing the oscillation phase and/or spatially structured excitability were performed by the newly introduced protocols as follows. Classically, the phase resetting has been performed by bath application of cAMP[28,31]. In our buffer-submerged culture condition, the application of cAMP (the final concentration of 20 and 50 nM for an early and late development, respectively) to the welldeveloped cell population synchronized the oscillation phase of the cAMP and allowed the reappearance of the symmetric circular waves but not the spiral waves (**Supplementary Fig S3a**, top). Such phase synchronization was simply recapitulated by the gentle mixing of the culture-buffer (**Supplementary Fig. S3a**, bottom). In detail, the cell population with the mature spirals were forced to be excited (**Supplementary Fig. S3a**, the first and second columns). Then the cell population synchronously fell into the refractory state (**Supplementary Fig. S3a**, the third column). Subsequent dynamics including the reappearance of the concentric waves (**Supplementary Fig. S3a,** the fourth column) and temporal synchronization of the local oscillation phase (**Supplementary Fig. S3b**) indicated that the simple mixing of culture-buffer homogenized the spatial distribution of the extracellular cAMP released at the wavefront, that eventually mimicked the bath application of cAMP.

For the spatial resetting, we performed an extensive mixing of the culture-buffer to detach cells from the surface of the culture dish. A few minutes after the dissociation, cells re-attached to the bottom of the culture dish with the cellular position being randomized and re-started the wave dynamics of cAMP. Concomitantly, the buffer-mixing had the phase synchronization effects as described above, thus results for the perturbation experiments should be read as summarised in **Supplementary Fig. S3c**.

### Supplementary Figures S1–S4:

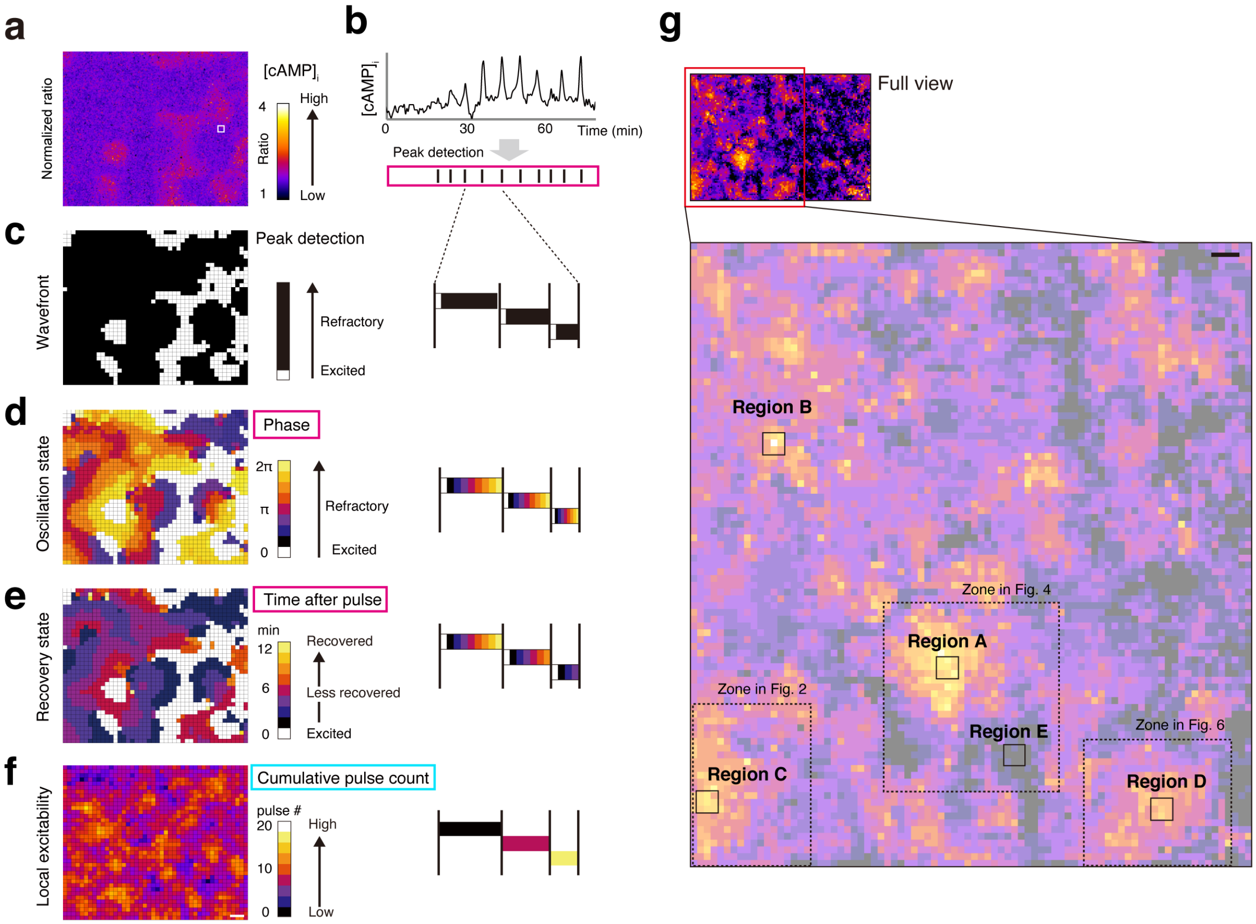


**Supplementary Figure S1. High-resolution Image Acquisition of Expanded FOV.**

(**a–e**) Data analysis scheme. Ratio image of mRFPmars/Flamindo2 (a). The cAMP signal in each analysis-ROI is subjected to peak detection (b). The binary representation of the pulse timing detects the wavefront (white is the excitation peak, c). Phase map constructed by the relative representation of the peak-to-peak interval (0 to 2π, d). Absolute representation of the elapsed time from the previous pulse as a measure of the recovery state (warmer color indicates greater recovery from the previous excitation (e). Cumulative pulse counts as an indicator of local excitability (f). (**g**) Location of areas presented in the main Figures. Scale Bars, 0.5 mm.

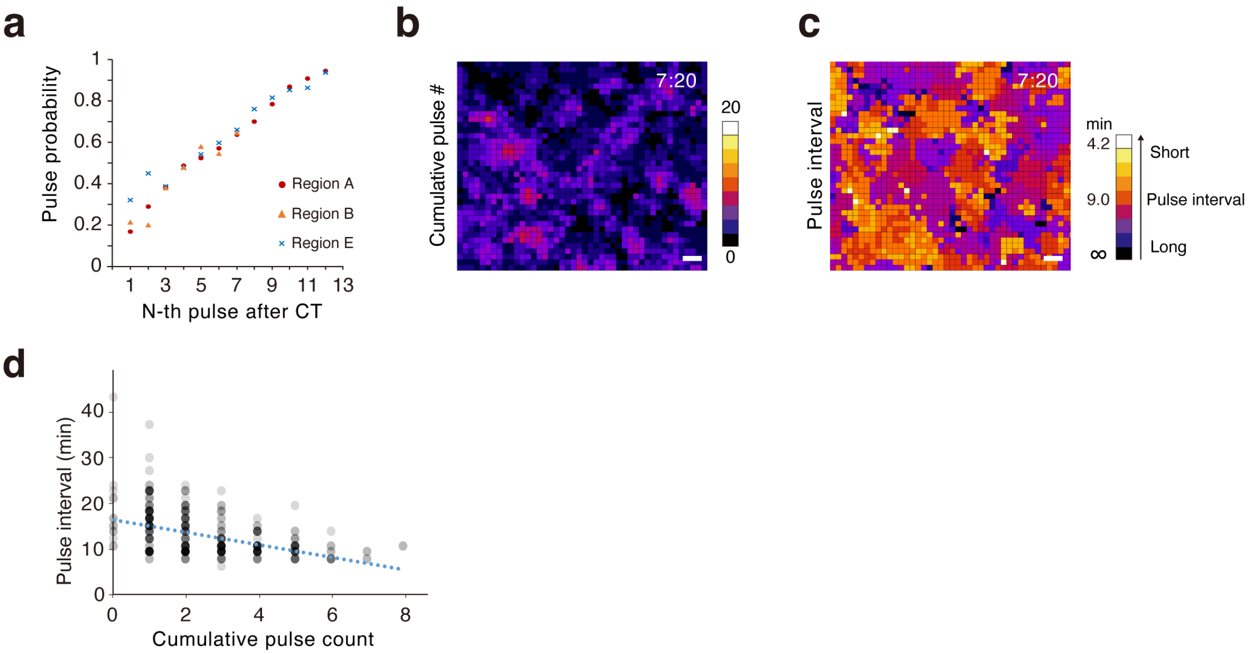


**Supplementary Figure S2. Pulse-dependent Increase in the Local Excitability.**

(**a**) Pulse probability in the post-CT period plotted against the population pulse in three areas with a high to low excitability (regions A, B, and E in **Supplementary Fig. S1g**). The conserved growth rate of the pulse probability in which a half-maximal pulse probability is yielded with a pulse count ~5. (**b-d**) Pulse-dependent increase in the oscillation frequency. The cumulative number of pulses (b) and local frequency (c) at 8 hours of the development. Correlation between the pulse count and local frequency in 1,672 ROIs (d). Scale Bars, 0.5 mm. See also **Supplementary Note S1**.

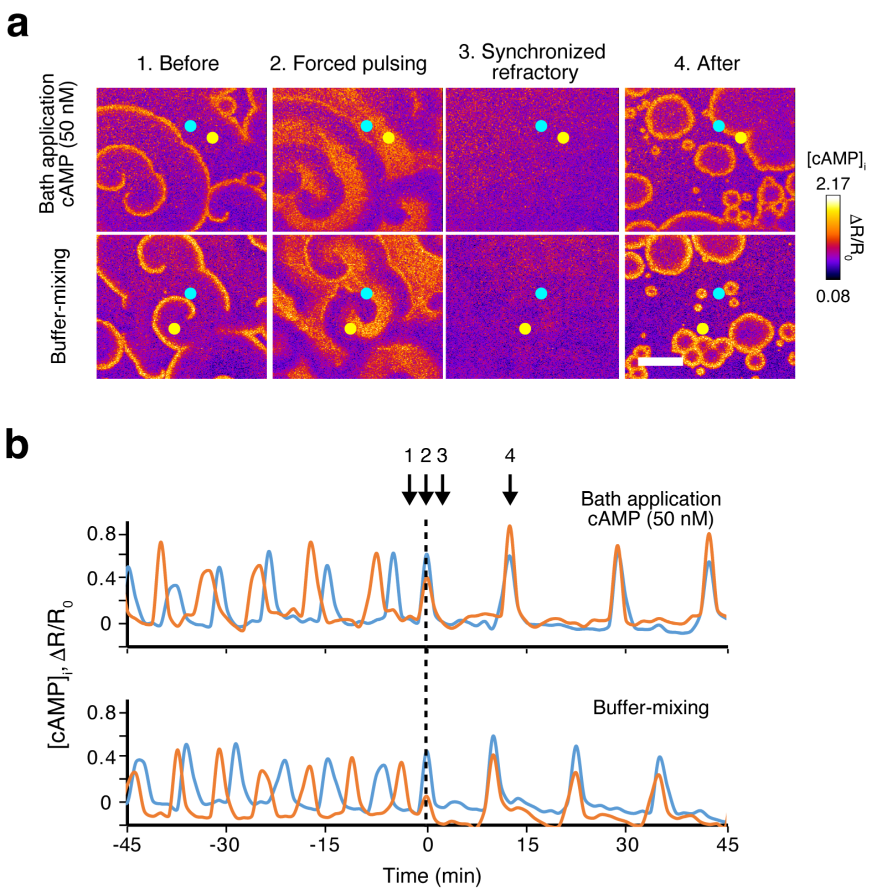


**c**

| Reset protocol (Name) | Phase reset | Spatial reset |
|---|---|---|
| Mixing type | Buffer (gentle) | Buffer & cell (intensitve) |
| Oscillation phase | Synchronized | Synchronized |
| Spatial excitability | Not affected | Randomised |

**Supplementary Figure S3. Wave Perturbation Experiments.**
(**a**) The phase resetting to the cell population having mature spiral waves by a bath application of cAMP (final 50 nM, top) and by the gentle buffer-mixing (bottom). Ratio images for $[cAMP]_{cell}$ at the pre-resetting (1), forced pulsing by the reset (2), synchronized refractory state (3), and the re-appeared cAMP wave (4). (**b**) The temporal synchronization of the oscillation phase in two ROIs (cyan and orange circles in a). (**c**) Summarised effects of the reset protocols. Scale Bars, 5 mm. See also **Supplementary Note S2**.

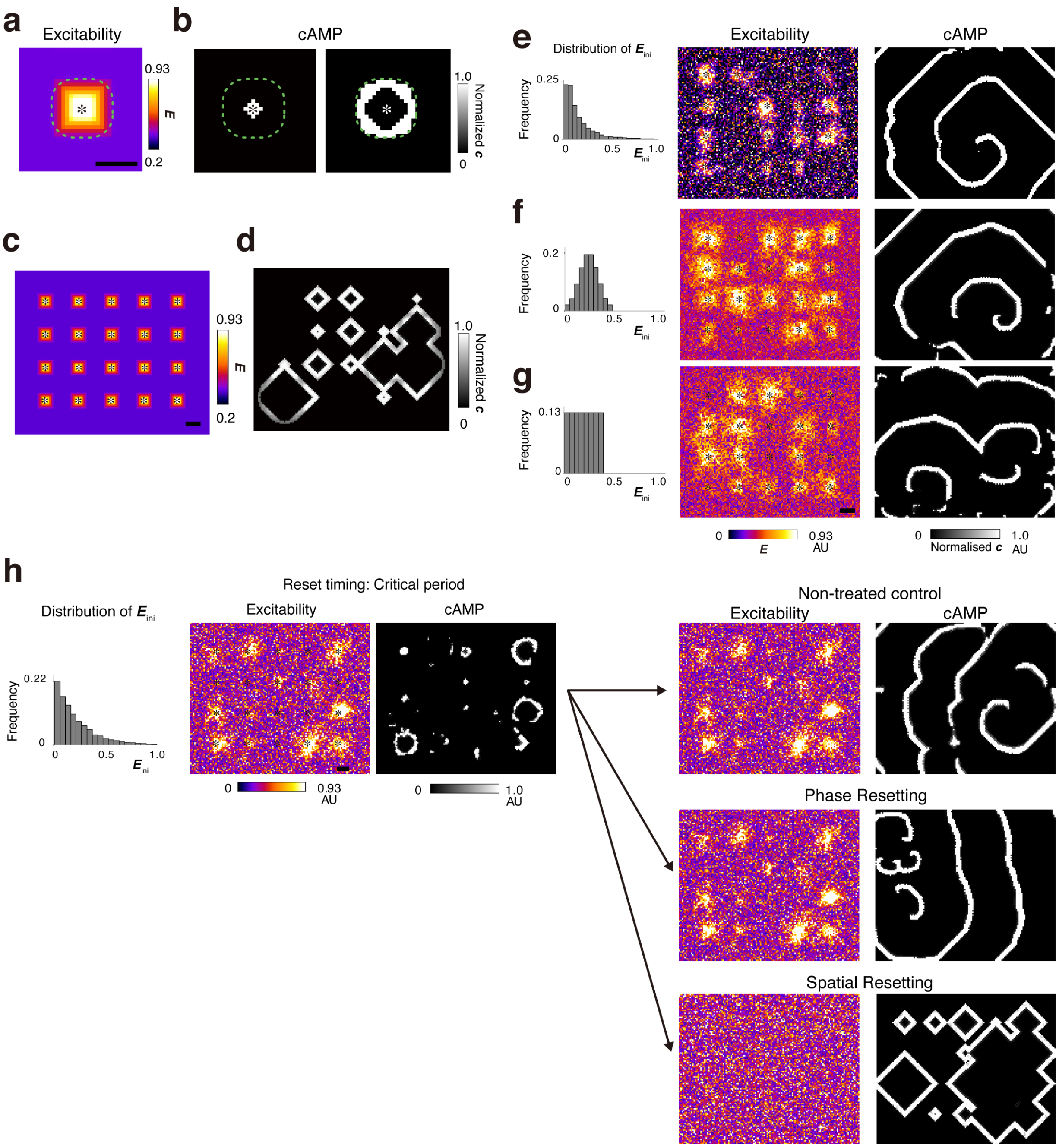


**Supplementary Figure S4. Modified CA Simulation Considering the Few PM Activities and Distributed Initial Excitability.**

(**a-d**) CA simulation for the homogeneous $E_{\mathrm{ini}}$. Isotropic growth of the high-excitability territory (dashed green line) seeded by a fixed pacemaker (asterisk) (a). Fragmentation of the cAMP wave does not develop spontaneously for the symmetric configuration of the high-excitability territory and the PM activity (b). Variation in the size of high-excitability territories and the wave fragmentation is not observed in a large-scale simulation (c, d). (**e-g**) Simulation for the distributed $E_{\mathrm{ini}}$, log-normal (e), Gaussian (f), and uniform distributions (g). The initial condition of $E$ (average value, 0.2, left), developed patterns of $E$ from the $4 \times 5$ array of the PM activity (middle) and cAMP wave (right). Asterisks are pacemakers. (**h**) Simulated effects of the phase and spatial resetting. Snapshots of *E* and *c* for non-treated control (top), phase (middle), and spatial resetting (bottom). The initial condition of *E* (Logarithmic distribution) is shown at the left. These results were replicated on three different data sets of $E_{\mathrm{ini}}$. Scale Bars, 0.5 mm.
